# Delineating the Organization of Projection Neuron Subsets with Multi-fluorescent Rabies Virus Tracing Tool

**DOI:** 10.1101/2019.12.16.877258

**Authors:** Liang Li, Yajie Tang, Leqiang Sun, Jinsong Yu, Hui Gong, Hannah C. Webber, Xiaoyu Zhang, Zhe Hu, Xiangning Li, Khaista Rahman, Zhenfang Fu, Jinxia Dai, Gang Cao

## Abstract

The elegant functions of the brain are facilitated by sophisticated connections between neurons, the architecture of which is frequently characterized by one nucleus connecting to multiple targets via projection neurons. Delineating the sub-nucleus fine architecture of projection neurons in a certain nucleus could greatly facilitate its circuit, computational, and functional resolution. Here, we developed multi-fluorescent rabies virus to delineate the fine organization of corticothalamic projection neuron subsets in the primary visual cortex (V1). By simultaneously labeling multiple distinct subsets of corticothalamic projection neurons in V1 from their target nuclei in thalamus (dLGN, LP, LD), we observed that V1-dLGN corticothalamic neurons were densely concentrated in layer VI, except for several sparsely scattered neurons in layer V, while V1-LP and V1-LD corticothalamic neurons were localized to both layers V and VI. Meanwhile, we observed a fraction of V1 corticothalamic neurons targeting multiple thalamic nuclei, which was further confirmed by fMOST whole-brain imaging. We further conceptually proposed an upgraded sub-nucleus tracing system with higher throughput (21 subsets) for more complex architectural tracing. The multi-fluorescent RV tracing tool can be extensively applied to resolve architecture of projection neuron subsets, with a strong potential to delineate the computational and functional organization of these nuclei.

## Introduction

The mammalian brain is a highly sophisticated networks of a vast number of neurons connected through specific neural circuits that have been roughly mapped for their contribution to certain brain functions such as sensation, movement control, learning and memory, emotion, wakefulness and sleep ^[1–8]^. Resolving the fine architecture and function of neural circuits is pivotal for understanding complex brain functions. Deciphering brain connections has been largely dependent on effective circuit tracing, measuring, and manipulation techniques. Neurotropic virus-based tracers have been widely utilized to elucidate the organization and function of neural circuits owing to their stronger signal, selectivity, compatibility with exogenous genetic engineering methods, and trans-synaptic ability ^[9–12]^. The insertion of exogenous genes such as optogenetic, chemogenetic, and Ca^2+^/voltage-sensitive genes in combination with a Cre/lox system allows for tracer viruses to be used for the monitoring, manipulation, and subsequent functional dissection, of specific neural circuits ^[13–16]^. Among various viral tracers, the rabies virus ^[17]^ has been widely used for direct input and mono-trans-synaptic upstream tracking, activity monitoring and manipulation of specific sets of labeled neurons ^[14, 18–24]^.

In the brain, single nuclei or functional brain regions (such as functional regions in the cerebral cortex, amygdala, and the nucleus of monoaminergic neurons) generally send outputs to multiple brain regions via projection neurons, while simultaneously receiving multiple inputs^[25–29]^. For example, the primary visual cortex (V1) projects to multiple thalamic nuclei including the dorsal lateral geniculate nucleus (dLGN), lateral posterior nucleus of thalamus (LP), and lateral dorsal nucleus of thalamus^[30]^ through reciprocal projections in which the V1-dLGN circuit was found to modulate thalamic responses to incoming visual signals and retinogeniculate targeting and refinement ^[31–33]^. Delineating the detailed organization of projection neuron subsets in a certain nucleus, or the fine architecture of projection neurons at a sub-nucleus level, is essential to the computing and functional resolving of circuits. Thus, a circuit tracing system with sub-nucleus resolution will facilitate the understanding of the organization and function of neural circuits stemming from the same brain region and extending out to multiple target regions.

In this study, we developed multi-fluorescent recombinant rabies viruses to tease out the sub-nucleus architectural organization of projection neuron subsets in certain brain nuclei. By simultaneously injecting multi-fluorescent RVs in thalamic targets of V1 projection neurons, we successfully mapped out the fine organization of V1 corticothalamic projection neuron subsets (V1-dLGN, V1-LP, and V1-LD), which is critical to better understanding the function of different corticothalamic projections in visual perception and vision-evoked behavior. Furthermore, we proposed a higher throughput (21 subsets) version of sub-nucleus tracing which will contribute to resolving the more subtle organizational features and functions of neural circuits throughout the nervous system.

## Materials and Methods

### Animals

Wild-type C57BL/6 mice were housed under standard conditions with food and water available *ad libitum*. Mice were maintained on a 12-h light/dark cycle at a temperature of 22–25°C. All the animal care procedures and experiments were approved by the Research Ethics Committee, Huazhong Agricultural University, Hubei, China (HZAUMO-2016-021), and were carried out in accordance with the Guide for the Care and Use of Laboratory Animals from Research Ethics Committee, Huazhong Agricultural University.

### Virus Construction and Preparation

To construct the multi-fluorescent RV tracing viruses, fluorescent protein genes (eGFP, tagBFP, or DsRed2) were cloned into multiple cloning sites of a pSAD-∆G-F3 backbone (a kind gift from Dr. Fuqiang Xu from Wuhan Institute of Physics and Mathematics, Chinese Academy of Sciences, and Dr. Edward Callaway from SALK Institute, USA) between Asc I and Sac II sites to generate pSAD-ΔG-tagBFP, pSAD-ΔG-eGFP, pSAD-ΔG-DsRed2 plasmids. Three recombinant RVs (rRV-ΔG-tagBFP, rRV-ΔG-eGFP, and rRV-ΔG-DsRed2) were rescued according to the Callaway laboratory’s protocol^[20, 34]^. In detail, 6 μg pSAD-∆G-F3 (Addgene, 32634) with different fluorescent protein genes and helper plasmids (3 μg pcDNA-SADB19N (Addgene, 32630), 1.5 μg pcDNA-SADB19P (Addgene, 32631), 1.2 μg pcDNA-SADB19G (Addgene, 32633) and 1.5μg pcDNA-SADB19L (Addgene, 32632) were co-transfected into 6-well culture plate with 10^6^ B7GG cells (kind gifts from Drs. Xu and Callaway) by FuGENE HD transfection reagent. The transfected B7GG cells were cultured for 7-10 days. Then the viral supernatant was harvested and spun at 4000 g for 15 min at 4°C to remove any debris. We then filtered the supernatant with a 0.45 μm filter (Millipore, SLHV033RS) and centrifuged the filtered sample at 70,000 g for 3 h, after which the pellet was re-suspended in 500 μL DMEM and added with 5 μL DNase I solution (Takara, M0303L) for a 30 min incubation at room temperature to avoid the viral particles aggregation. We loaded the virus-DMEM-DNase I mixture on the top of 20% sucrose solution (Sinopharm Chemical Reagent Co., Ltd, 10021418) and spun it again to purify the virus. Finally, rRVs were re-suspended in PBS, aliquoted and stored at −80°C. The rRV titration was performed with infection of BSR cell line seeded in 96-well plate by 10x gradient dilution of rRV. The titer of the rRVs was adjusted to 10^9^ plaque-forming units /mL in preparation for their injection into the mouse brains.

### Construction of Plasmids for Subcellular Signal Localization

The subcellular localization sequences, including the nuclear localization sequence (NLS, 5’-CCTCCAAAAAAGAAGAGAAAGGTC-3’), mitochondrial localization sequence (Mito, 5’-ATGCTTTCACTACGTCAATCTATAAGATTTTTCAAGCCAGCCACAAGAA CTTTG-3’), and cell membrane localization sequence (Mem, 5’-AAGCTGAACCCTCCTGATGAGAGTGGCCCCGGCTGCATGAGCTGCAAGTGTGTGCTCTCC-3’), were fused to the end of the EGFP gene. The fused genes were then inserted into the eukaryotic expression vector PCAGGS between EcoR I and Bgl II enzyme sites to generate PCAGGS-EGFP-NLS, PCAGGS-EGFP-Mito, and PCAGGS-EGFP-Mem plasmids. These plasmids were transfected along with Lipofectamine 2000 transfection reagent (Thermo Fisher Scientific, USA) into HEK293 T cells. These cells were later fixed with 4% paraformaldehyde, counter stained with DAPI, and imaged by confocal microscopy (Olympus, Japan) to confirm the subcellular localization of the EGFP fusion proteins.

### Stereotaxic Microinjection

Stereotaxic microinjection was performed according to the protocol by Peter H Seeburg^[35]^. Male C57BL/6 mice (8-10 week-old) were anesthetized by intraperitoneal injection of mixed anesthetics (7.5 g urethane, 3 g chloral hydrate, and 75 mg xylazine in 100 mL ddH_2_O) at 9 μL/g body weight. The scalps of the mice were shaved and cleaned with 70% ethanol. The heads of the mice were then fastened in the stereotaxic instruments (68025; RWD Life Science, Shenzhen, China). Holes were drilled into the skull using a 0.5 mm diameter microdrill (78001; RWD Life Science) at the sites of intended virus injection into the dorsal lateral geniculate nucleus (dLGN; AP, –2.35 mm; ML, 2.26 mm; DV, –2.70 mm), the lateral posterior nucleus (LP; AP, –2.22 mm; ML, 1.60 mm; DV, –2.62 mm), and the lateral dorsal nucleus (LD; AP, – 1.32 mm; ML, 1.40 mm; DV, –2.65 mm). The injection apparatus consisted of a 10 μL syringe (Shanghai Gaoge Industrial and Trading Co., Shanghai, China) filled with mineral oil and fused with a glass needle pulled by PC-10 puller (DL Nature gene Life Sciences, Newbury Park, CA). The virus was injected using a KDS Legato™ 130 micro-pump (KD Scientific Inc., Holliston, MA) at a speed of 18 nL/min, with a total injection volume of 300 nL virus (10^9^ focus-forming units/mL).

### Brain Slice Preparation, Imaging and Reconstruction

Mice were anesthetized and perfused with 4% paraformaldehyde in 0.01M PBS. Mouse brains were dissected out and post-fixed in 4% paraformaldehyde in 0.01M PBS overnight at 4°C, then embedded in 5% agarose gel and sectioned into 40 μm slices using a 7000SMZ vibratome (Campden Instruments, United Kingdom). These slices were mounted on glass slides and sealed with mounting oil (enamel from cosmetics shop). Mounted brain slices were imaged using a BX63 fluorescent microscope (Olympus, Japan) or a laser scanning confocal microscope (Olympus, Japan).

### fMOST Imaging and Single Neuron Tracking

Fluorescence micro-optical sectioning tomography (fMOST) imaging of single neurons in the whole brain wide was performed as previously described^[36–38]^. The fMOST imaging system consists of brain tissue preparation and embedding with resin, automatic sectioning, continuous data acquisition, raw images modulation, and reconstruction, as well as single neuron tracking using specialized software^[36]^. Firstly, the whole brains of mice were dissected out after transcardial perfusion with 4% paraformaldehyde in 0.01M PBS, and post-fixed in 4% paraformaldehyde in 0.01M PBS overnight at 4°C. Then, mice brains were rinsed in 0.01M PBS solution (50 mL per mouse brain) overnight and subsequently dehydrated in a graded ethanol series (50%, 75%, 95%, and three times of 100%) for 2 h each time at 4°C. The dehydrated brains were impregnated in graded (70%, 85%, and 100%) solution of Glycol Methacrylate water-soluble resin (GMA solution, Ted Pella Inc., Redding, CA) for 2 h at each immersion time, followed by an overnight immersion in a 100% GMA solution and subsequent immersion in pre-polymerized GMA for 3 days at 4°C. The whole brain was then embedded in a gelatin capsule pre-filled with pre-polymerized GMA and polymerized for 60 h at 60°C. To improve GFP fluorescence signal, 1 M NaOH was added to each of the GMA infiltrate solutions to reach 0.3% volume. Then, the whole brain was sectioned into ultra-thin slices (2 μm thick) using a diamond knife with a three-dimensional nano-precision translation stage and imaged continuously at a voxel size of 0.32×0.32×2 μm^3^ for the generation of high-resolution data (total 5000 high-resolution images per mouse brain). Each time, one small 3D data of a ~0.125 mm^3^ block of the whole brain was loaded into Amira 5.4.1 software (FEI, Mérignac Cedex, France) for neuron tracking analysis. Entire outlines of single neurons were manually traced from axon initiation to boutons by combing data blocks using Amira 5.4.1 software.

### Image Analysis

Serial images of the whole brain were aligned to create a 3-D construction using Auto aligner software (Bitplane AG, Switzerland), then rendered and exported to a 3-D view video or image format using Imaris 8.0 (Bitplane AG, Switzerland), as shown in figure 2N.

## Results

### Development of Multi-fluorescent Rabies virus

In order to tease apart the fine architecture of certain brain nuclei at sub-nucleus resolution based on their various projection targets, we developed multi-fluorescent RV tracing viruses. Distinct projection targets of a specific brain nucleus can be simultaneously injected with recombinant rabies viruses (rRV) expressing different fluorescent proteins which can retrogradely label the nucleus from its targeted brain regions, thus tracing the fine architecture of different projection neuron subsets (Fig.1A). Firstly, the recombinant rabies viruses (rRV) rRV-ΔG-tagBFP, rRV-ΔG-eGFP, and rRV-ΔG-DsRed2 were engineered using reverse genetics, by which the glycoprotein was replaced with tagBFP, eGFP, or DsRed2 genes, respectively, to incorporate the fluorescent labeling strategy and avoid trans-synaptic spreading of the RV (Fig.1B). In this scenario, the rRV tracing virus can retrogradely labels projection neurons by transducing the axon terminal at target sites, as shown by the specific labeling of V1 corticothalamic projection neurons from their thalamic targets (dLGN, LP or LD) in Fig.1C1–1E3. The mean number of labeled projection neurons by rRV tracing system in the whole V1 cortex is 7773, 15882 and 18096 neurons for V1-LD, V1-LP and V1-dLGN respectively (n=3 mice for each injection site). Strong fluorescent labeling of rRV infected neurons revealed fine neuronal structure, including resolution of axons and dendrites (Fig.1C1–1E3). Most of the V1-dLGN corticothalamic neuron somas are densely located in layer VI except for several sparsely scattered neurons in layer V along the entire V1 cortex. In contrast, the somas of V1-LP and V1-LD corticothalamic neurons were mainly located in layers V and VI of the V1 cortex (Fig.1C1–1E3).

**Figure 1.**
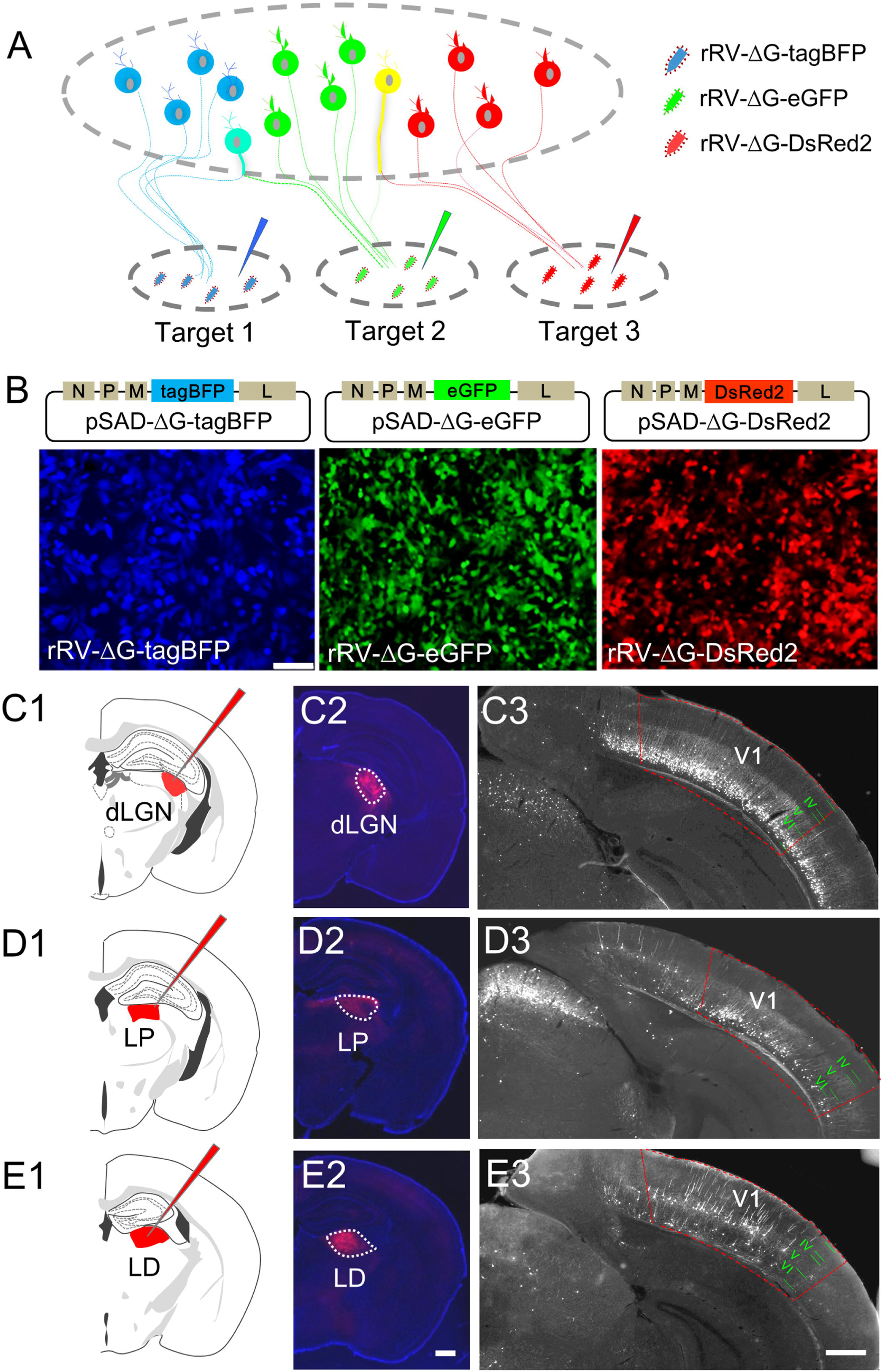
Development of multi-fluorescent RV tracing virus. **(A)** Diagram of sub-nucleus delineation of projection neuron subsets in certain brain nucleus by recombinant rabies virus (rRV) with different fluorescent protein genes (rRV-ΔG-tagBFP, rRV-ΔG-eGFP, and rRV-ΔG-DsRed2). **(B)** Construction and preparation of rRV-ΔG-tagBFP, rRV-ΔG-eGFP, and rRV-ΔG-DsRed2 tracing viruses. TagBFP, eGFP and DsRed2 genes were cloned into RV pSAD-∆G backbones, respectively, and the recombinant RVs (rRV-ΔG-tagBFP, rRV-ΔG-eGFP, and rRV-ΔG-DsRed2) were successfully rescued in B7GG cells as indicated by fluorescent protein expression. Scale bar, 100 μm in B. **(C1-E3)** Tracing viruses of rRV-∆G-eGFP were injected into dLGN, LP and LD, respectively, as indicated by C1-E1. The accuracy of injection sites in thalamic nuclei was confirmed by CTB-Alexa594 tracer injection, as displayed in C2-E2. Bright fluorescence in the cell body and dendrites, allowing for the fine labeling of neurons, was observed in retrogradely labeled projection neurons in V1. The V1 cortex is outlined by red frames. Scale bars, 500 μm in C2-E2 and C3-E3. rRV, recombinant rabies virus; ∆G, deleted glycoprotein; V1, primary visual cortex; LD, lateral dorsal nucleus of thalamus; LP, lateral posterior nucleus of thalamus; dLGN, dorsal lateral geniculate nucleus of thalamus; IV, layer 4 of V1 cortex; V, layer 5 of V1 cortex; VI, layer 6 of V1 cortex.

### Sub-nucleus Delineation of V1 Corticothalamic Neuron Subsets by Multi-fluorescent Rabies virus

To map the spatial organization of different corticothalamic neuron subsets in the same V1 cortex, three rRVs (rRV-ΔG-tagBFP, rRV-ΔG-eGFP, and rRV-ΔG-DsRed2) were simultaneously injected into thalamic targets of V1 (dLGN, LP, and LD) (Fig.2A). Consistent with the results of Fig.1, V1 corticothalamic neuron subsets with distinct fluorescent colors were distributed throughout layers V and VI, though largely non-overlapping and in clearly distinct patterns (Figure 2B–2N). The V1-dLGN corticothalamic neurons are preferentially localized to layer VI of the V1 cortex (Fig.2B-2D, 2N). In contrast, V1-LP corticothalamic neurons are concentrated in layers V and VI of the V1 cortex, showing a clear separation between the two distribution layers (Fig.2E-2G, 2N). Similarly, V1-LD corticothalamic neurons were mainly located in layers V and VI along the entire V1 cortex (Fig.2H-2J, 2N). Furthermore, the serial brain sections were reconstructed into 3-D images to demonstrate the spatial organization of three subsets of V1 corticothalamic neurons in coronal views (Fig.2K-2M, 2N). We show that V1-dLGN corticothalamic neurons are largely concentrated in layer VI of V1, whereas V1-LP and V1-LD corticothalamic neurons are distinct and located in both layers V and VI.

**Figure 2.**
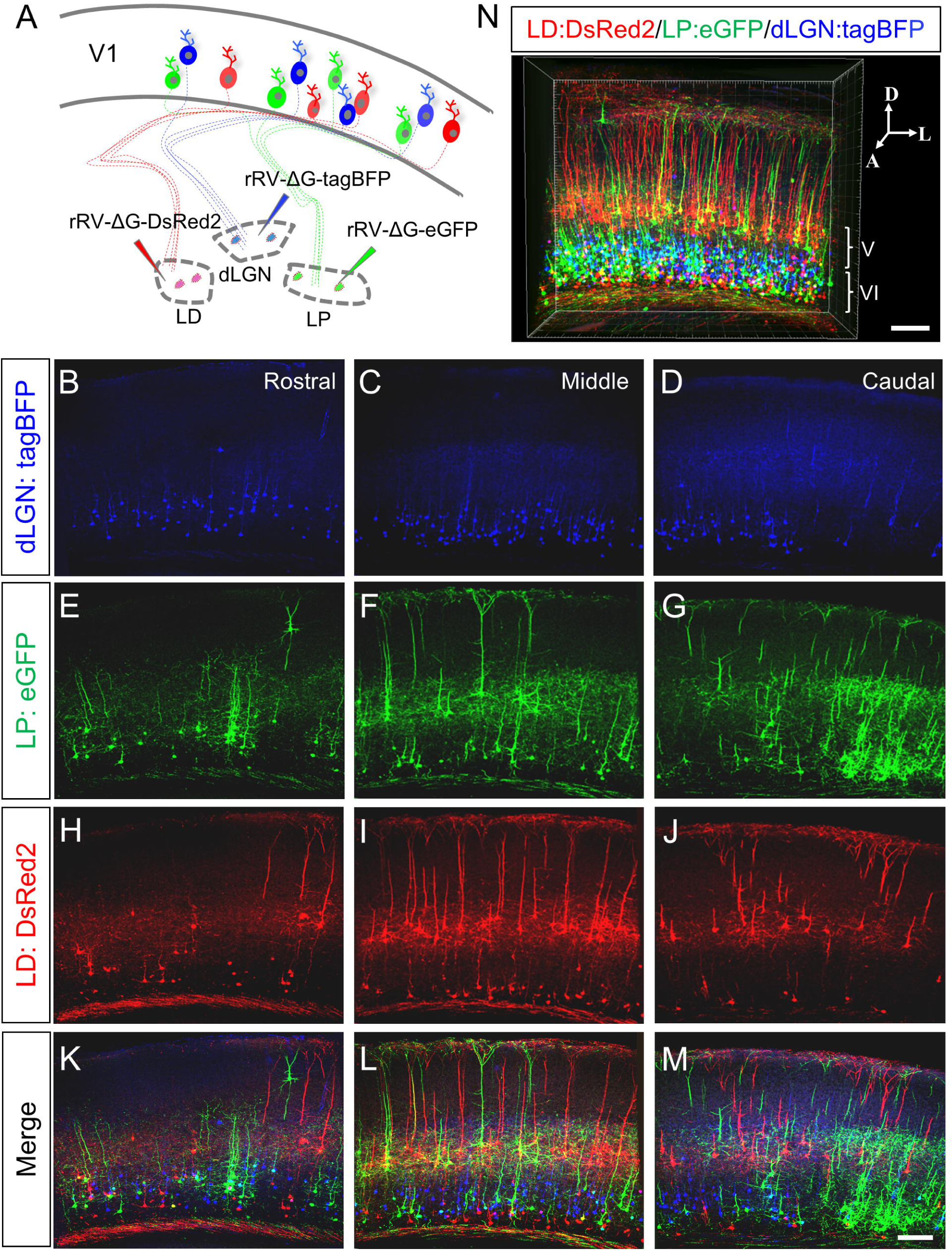
Resolving the architecture of V1 corticothalamic neuron subsets using multi-fluorescent RV. **(A)** Schematic illustration of retrograde labeling of V1 corticothalamic neuron subsets by simultaneous injection of rRV-ΔG-tagBFP, rRV-ΔG-eGFP, and rRV-ΔG-DsRed2 into dLGN, LP and LD respectively. **(B-M)** V1 corticothalamic neuron subsets (V1-dLGN, V1-LP, and V1-LD) were retrogradely labeled as shown in separate and merged images of coronal sections in the rostral, middle, and caudal V1 cortex. Scale bars, 150 μm in B-M. **(N)** The continuous coronal brain images were reconstructed into a 3-D image. White brackets mark layers V and VI of V1. Scale bars, 200 μm in N. V1, primary visual cortex; LD, lateral dorsal nucleus of thalamus; LP, lateral posterior nucleus of thalamus; dLGN, dorsal lateral geniculate nucleus of thalamus.

During the triple-site retrograde labeling of V1 corticothalamic neuron subsets, a small fraction of V1 corticothalamic neurons were observed to be co-labeled with two different fluorescent tracers (Figure 3), indicating that these V1 corticothalamic neurons simultaneously project to multiple thalamic nuclei. As shown in Figure 3A, GFP-positive V1-LP corticothalamic neurons (as indicated by arrows) are co-labeled with DsRed2 fluorescence, indicating that they also project to LD. In figure 3B, tagBFP-positive V1-dLGN corticothalamic neurons (as indicated by dashed circles) were co-localized with eGFP (V1-LP corticothalamic neurons). The dashed box indicated tagBFP-positive V1-dLGN projection neurons co-labeled with DsRed2 (V1-LD corticothalamic neurons).

**Figure 3.**
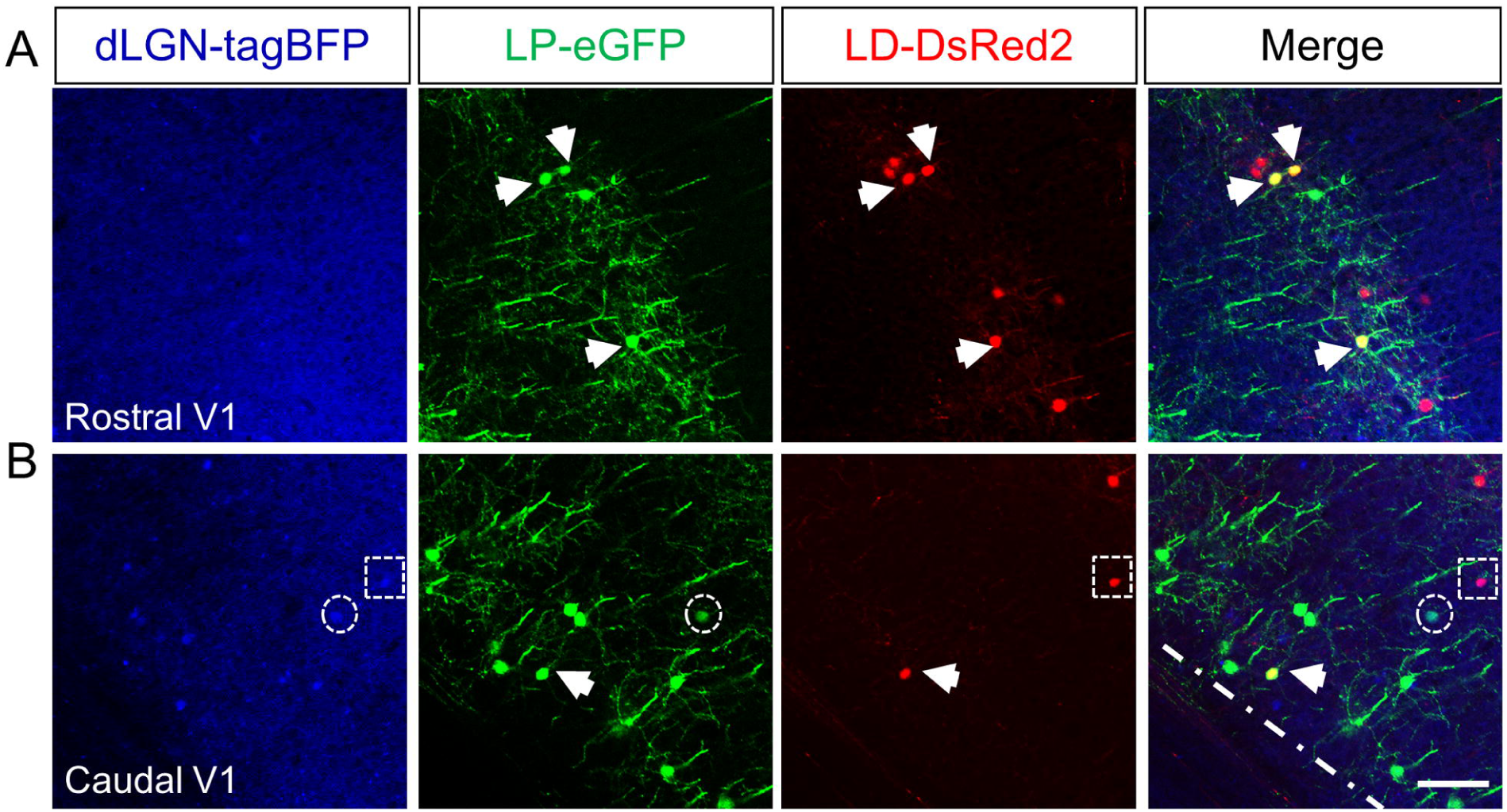
Labeling of multi-target projection neurons in V1. After simultaneous injection of rRV-ΔG-tagBFP, rRV-ΔG-eGFP, and rRV-ΔG-DsRed2 in dLGN, LP and LD respectively, V1 multi-target projection neurons were observed by overlapping of different colors as indicated by white arrows for LP/LD targeting projection neurons, dashed circles for dLGN/LP targeting projection neurons and dashed boxes for dLGN/LD targeting projection neurons in both rostral **(A)** and caudal **(B)** V1. Scale bars, 100 μm. V1, primary visual cortex; LD, lateral dorsal nucleus of thalamus; LP, lateral posterior nucleus of thalamus; dLGN, dorsal lateral geniculate nucleus of thalamus.

To further obtain an overall view of these V1 multi-target projection neurons, fluorescence micro-optical sectioning tomography (fMOST) imaging was applied to track the entire axon of a single V1 corticothalamic neuron through the entire brain region (Figure 4A). To this end, rRV-ΔG-eGFP was injected into the dLGN and the brain was fixed with resin for fMOST imaging. The detailed morphology of the retrogradely labeled V1 corticothalamic neurons was reconstructed by fMOST as shown in figure 4B. Three randomly selected corticothalamic neurons with multiple targets in V1 along anterior-to-posterior sections were traced manually throughout the whole brain, the precise projection sketch of which is displayed in figure 4C-4E. The fMOST imaging results clearly demonstrate that a fraction of V1-dLGN corticothalamic neurons can indeed simultaneously project to different thalamic nuclei (dLGN, LP, LD) and even other brain regions including SC and V2L through collateralization (Figure 4C–4F), suggesting that such corticothalamic neurons in V1 can modulate multiple nuclei including the visual thalamic nucleus (dLGN, LP, LD), SC, and V2. A primary fMOST 2D image of coronal brain section is shown in Figure S1.

**Figure 4.**
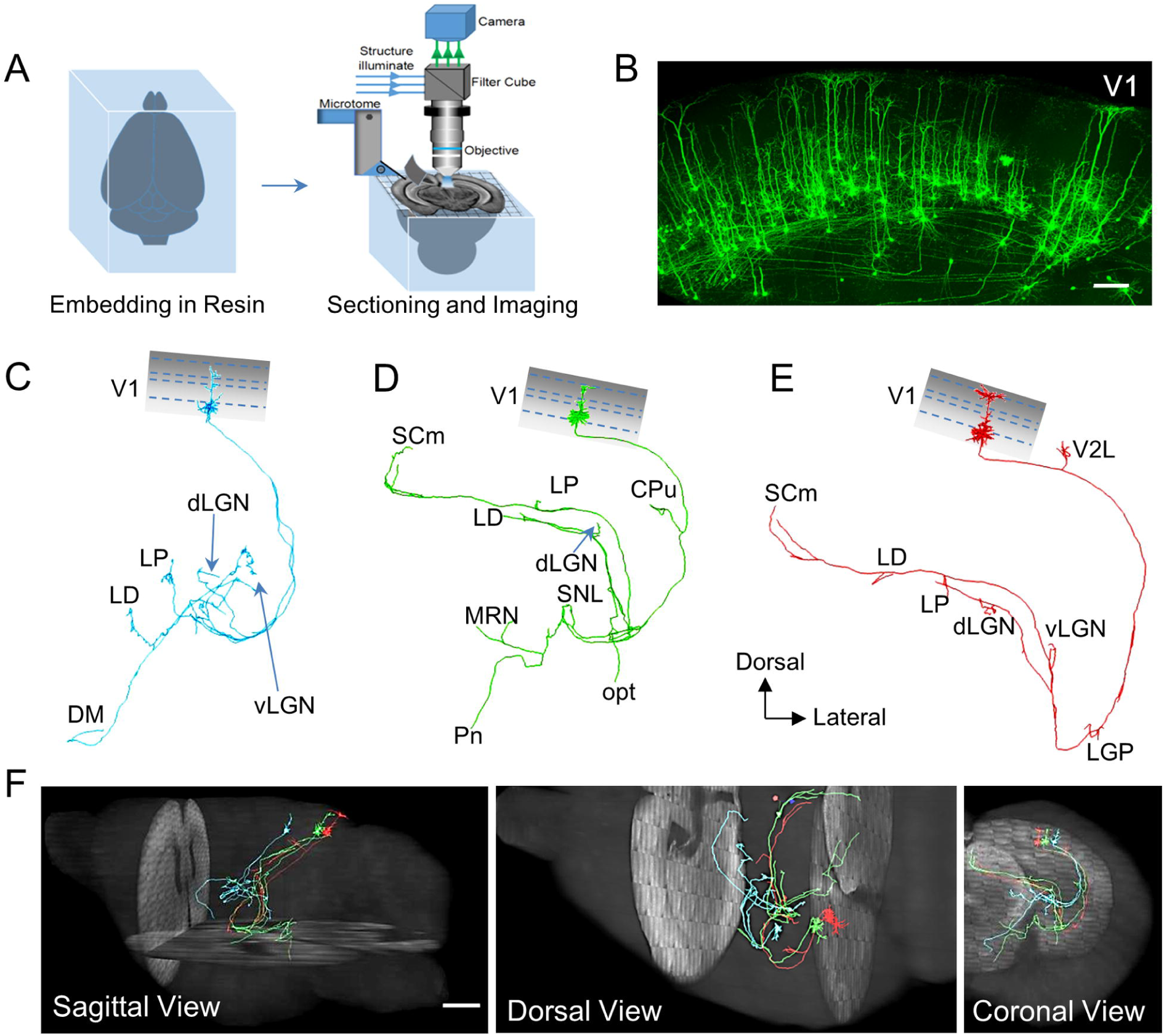
Tracking whole-brain projections of V1 multi-target corticothalamic neurons by fMOST imaging. **(A)** Schematic illustration of fMOST imaging. Mouse brains embedded in resin were sectioned using a microtome and simultaneously imaged by laser confocal microscopy. Finally, the continuous images of the whole brain were reconstructed for 3-D visualization. **(B)** The morphological details of the dLGN-targeting of V1 projection neurons labeled by rRV-∆G-eGFP were reconstructed by fMOST imaging. In total, 40 continuous brain slices of fMOST images were reconstructed to create a 3D image with a thickness of 80 μm. Scale bar, 200 μm in B. **(C-E)** The projection sketches of three randomly selected V1 multi-target projection neurons were manually traced from fMOST images. These neurons were retrogradely labeled by rRV-∆G-eGFP from the dLGN and displayed multi-target projection patterns throughout the whole brain. **(F)** A 3-D view of the whole brain projection sketch of V1 multi-target projection neurons by fMOST imaging (See also: movie S1). Scale bar, 1 mm in F. V1, primary visual cortex; V2L, lateral area of secondary visual cortex; LD, lateral dorsal nucleus of thalamus; LP, lateral posterior nucleus of thalamus; dLGN, dorsal lateral geniculate nucleus of thalamus; vLGN, ventral lateral geniculate nucleus of thalamus; DM, dorsomedial hypothalamic nucleus; SCm, motor-related superior colliculus; CPu, caudate putamen (striatum); SNL, lateral part of substantia nigra; MRN, midbrain reticular nucleus; opt, optic tract; Pn, pontine nuclei; LGP, lateral globus pallidus.

## Discussion

Delineating the fine architecture of neural circuits is desperately required to dissect and elucidate their functions. Most brain nuclei send projections to multiple target regions, while the detailed organization of projection neuron subsets to different targets within the same nucleus remains largely elusive. Therefore, developing a more effective and applicable tracing technique for the precise delineation of neural circuit at higher, sub-nuclear resolution will contribute to our understanding of the brain’s connectivity patterns and related functions. Here, we constructed a sub-nucleus resolution tracing system using the rabies virus to delineate the fine organization of V1 corticothalamic neuron subsets (V1-dLGN, V1-LP, and V1-LD). Due to the retrograde tracing property and strong labeling signal of rRV, our methods can build circuit-based neuron subsets mapping, which is difficult to implement solely by mice genetics. We observed distinct organizational patterns of different V1 corticothalamic neuron subsets, characterized by dense V1-dLGN corticothalamic neurons in layer VI and a distribution of V1-LP and V1-LD corticothalamic neurons in layers V and VI. The sub-nucleus architecture of V1 corticothalamic projection neurons is important for computational and functional studies of different corticothalamic neuron subsets in V1 (V1-dLGN, V1-LP, and V1-LD). The distribution of different subsets in V1 cortex is consistent with previous reports in which similar RV-ΔG or retro-beads tracer were injected into dLGN and resulted in the labeling of layer 6 corticothalamic neurons of V1^[20, 39]^.

It is of great interest to investigate the mechanisms underlying the distribution pattern of different V1 corticothalamic neuron subsets. One possible mechanism may be that a group of specific neurons wire together and follow the same axon guidance cues during development. It would therefore be conceivable that neurons belonging to a specific subset might express some common signature genes. The transcriptome of LP- and dLGN-targeting V1 neurons has been studied, indeed identifying some circuit specific genes such as Tph2, Cdh13 and Chrna6^[30]^. Further investigation on gene expression profiles of different V1 corticothalamic neurons subsets during development will help inform the molecular mechanisms of V1 corticothalamic neuron subsets formation.

Of note, using multi-fluorescent RV tracing system alongside fMOST imaging, we observed that certain V1 corticothalamic neurons can simultaneously project to multiple targets including visual thalamic nuclei, SC and V2, implying that the transmission of information to distinct visual circuits during the computation and integration of visual signals may be synchronized. Our findings are directly relevant to future studies on the functions of multi-target projection neurons in the V1 cortex in combination with various electrical activity detection and manipulation techniques such as fiber photometry, optogenetics, and chemogenetics.

Besides RV tracing virus, other retrograde viral tracers (such as canine adeno virus type 2 (CAV2), retrograde adeno associated virus (retro-AAV)) or traditional tracers (CTB, Fluorogold) are also utilized for tracing projection neurons to certain target regions^[17, 40–43]^. Compared with traditional tracers, viral tracers are more widely used to study the fine structural and functional properties of neural circuits due to their stronger signal, selectivity, genetic engineering convenience, especially trans-synaptic ability for higher order circuit tracing^[9–12]^. With the supply of RV glycoprotein by AAV virus to rRV-ΔG infected neurons, upstream neurons can also be labeled due to the rRV trans-synapse spreading^[14,19,22,44]^. The CAV2 and retro-AAV cannot spread trans-synaptically, limiting their use in higher upstream tracing. Thus, combining our multi-fluorescence RV tracing system with RV glycoprotein supplement by AAV virus, direct upstream neurons to different projection neuron subsets of V1 corticothalamic neurons can be tracked and analyzed.

Our tracing tools may be extensively applied to trace specific nuclei with multiple targets in the brain for mapping of projection neuron subsets. For the resolution of nuclei with a higher number of projection targets (>3 to 21), we conceptually propose a strategy consisting of engineering an upgraded version of a multi-fluorescent RV toolkit. In this updated toolkit, dozens of fluorescent protein genes (CFP, eGFP, YFP, mOrange2, mPlum, mStrawberry and IFP1.4) can be fused with different subcellular localization signaling peptides (cell membrane, nucleus, and mitochondria) to create 21 fluorescent labeling codes (Figure 5). Since membrane, nucleus, and mitochondria labeling of HEK293 cells displayed very distinct patterns of expression (figure 5A), it is easy to recognize the subcellular labeling in neurons by epifluorescence microscopy or confocal microscopy. Labeling can also be differentiated according to excitation/emission wavelength for CFP (458/480 nm for tagCFP, 433/475 nm for eCFP), eGFP (488/507 nm), YFP (513/527 nm), mOrange2 (549/565 nm), mPlum (590/649 nm), mStrawberry (574/596 nm) and IFP1.4 (684/708 nm). Such higher throughput sub-nucleus tracing system would allow for the architectural mapping of more complicated projection neuron subsets in certain brain nuclei. However, the simultaneous injection of 21 viruses into different brain sites is very challenging, which demands high expertise in microinjection. Moreover, lesser injection volume (100-150 nL) of tracing virus should be considered for small target regions to avoid overlap of tracing virus when performing projection neuron subsets mapping. A smaller 100-150 nL volume of RV-∆G tracing virus is effective in retrograde labeling of projection neurons such as projection neuron targeting to lateral hypothalamic area or dLGN according to our previous reports^[45]^.

**Figure 5.**
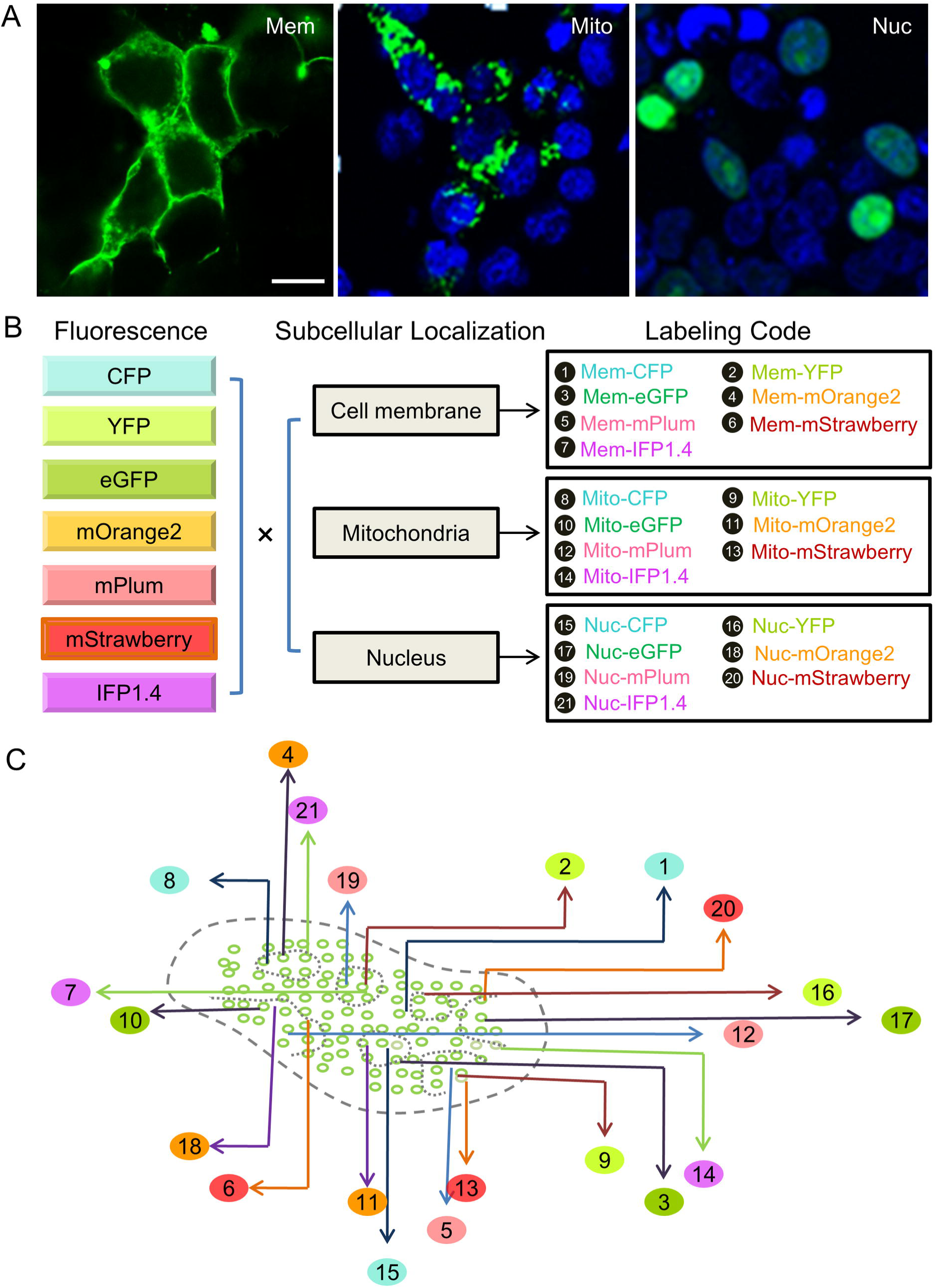
Proposed sub-nucleus tracing system of projection neurons with higher throughput (21 subsets). **(A)** Subcellular localization of fluorescent proteins can be assessed by fusion with a cellular organelle location or targeting sequence. The eGFP gene was fused with a location sequence for the cell membrane (Mem), mitochondria (Mito), or nucleus (Nuc), successfully allowing subcellular localization of eGFP protein following transfection in HEK293 T cells. Scale bar, 15 μm in A. **(B)** Combination of three subcellular locations with seven fluorescent proteins can generate 21 fluorescent labeling codes. **(C)** Using the upgraded version of the sub-nucleus tracing system based on the rRV system, the fine organization of larger numbers of projection neuron subsets (>3 to 21) in certain brain nuclei could be mapped at sub-nucleus resolution. **Movie S1.** 3-D reconstruction of three V1 multi-target projection neurons in the whole brain (also shown in Figure 4).

Moreover, recent studies showed that engineered RV strains and N2C strains of RV could increase tracing efficiency ^[54]^, indicating a method to further improve the current tracing strategy’s efficiency. Taken together, our multi-fluorescent RV retrograde labeling can be extensively applied to the tracing of the detailed architecture of any brain region with multiple projection targets, broadly contributing to the delineation of the brain’s fine circuits at sub-nucleus resolution.

## Supporting information

Movie S1. 3-D reconstruction of three V1 multi-target projection neurons in the whole brain (also shown in Figure 4).

## Acknowledgement

We thank Dr. Fuqiang Xu (Wuhan Institute of Physics and Mathematics, Chinese Academy of Sciences) and Dr. Edward Callaway (The SALK Institute, USA) for rRV packing system, and Dr. Xiaobing He and Ting Ding (Wuhan Institute of Physics and Mathematics, Chinese Academy of Sciences) for suggestions related to the RV preparation. This work was supported by the National Natural Science Foundation of China (Grant No.31700934, 31371106) and the Huazhong Agricultural University Scientific & Technological Self-innovation Foundation (Program No. 2662017PY082, 52204-13002, 2014BQ019).

---

**Figure S1.**
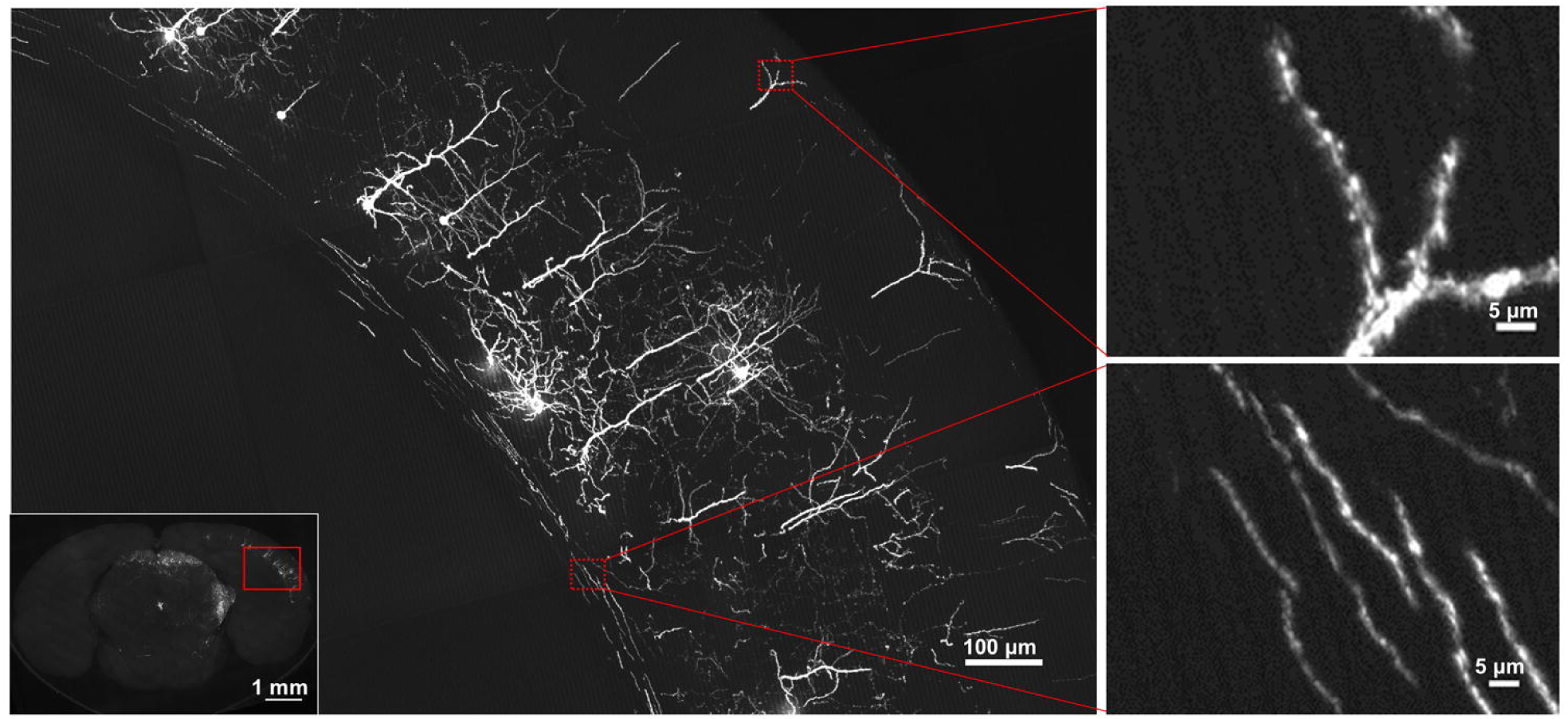
A primary fMOST image of mouse coronal brain section. The left panel is an enlarged view of outlined region (solid red rectangle) in the fMOST image of single coronal brain section (lower inset), with an image thickness of 2 μm. The top and bottom panels on the righ are high-magnification images of dendrites and axons respectively, which show the high resolution of single neuron tracing in the whole mouse brain. Resolution is 0.32×0.32×2 μm^3^ per voxel.

